# Evolutionary stability of microbial communities with antibiotic degrading species

**DOI:** 10.1101/045732

**Authors:** Eric D. Kelsic, Kalin Vetsigian, Roy Kishony

**Affiliations:** Department of Genetics, Harvard Medical School, Boston, Massachusetts; Wyss Institute for Biologically Inspired Engineering, Boston, Massachusetts, USA; Department of Bacteriology and Wisconsin Institute for Discovery, University of Wisconsin-Madison, Madison, Wisconsin, USA; Department of Systems Biology, Harvard Medical School, Boston, Massachusetts, USA; Faculty of Biology and Department of Computer Science, Technion-Israel Institute of Technology, Haifa, Israel

## Abstract

A major puzzle in ecology is how antibiotic resistant, sensitive and producer species coexist in close proximity. Recently, we showed that antibiotic degrading species dramatically alter community dynamics: replacing intrinsic resistance with resistance through degradation generates stable communities resilient to spatial mixing, large species abundance perturbations, changes in interaction strengths, and differences in species growth rates^1^. In addition to ecological stability, it is interesting to consider evolutionary stability of these communities to the appearance of cheater species that either cease production or degradation of antibiotics. Our investigation of evolutionary stability of cyclical 3-species communities revealed that these communities are robust to cheaters that stop degrading antibiotics. Our simulations also showed that cheaters that stop producing antibiotics do not take over the community^1^, yet they can transiently invade and cause community collapse^2^. In the analytical approximation we initially investigated, production cheaters with a small fitness advantage can invade the community simply because the benefit of inhibiting competitors is shared among all cells^2^. Here, we consider evolutionary stability to cheaters in our complete model^1^, where spatial mixing is introduced only after a short range colonization step. In this regime, an antibiotic producer cell directly benefits from killing nearby competitor species as it has a greater chance of colonizing the newly voided spaces created by the action of its antibiotic. Simulating response to cheater invasions for the cyclical three species community and for random 4-species ecologically stable topologies, we find that these communities can be fully resilient to both degradation and production cheaters. The strength of selection against cheaters varies with the area of the zone of inhibition around producers and is maximized for weak inhibition, where there is less overlap between the killing zones of neighboring cells. These results may aid the construction of complex synthetic communities that are both ecological and evolutionary stable.

---

Ecological models of antibiotics can lead to relationships of cyclic dominance among species (for example, rock–paper–scissors games), which can support coexistence in spatial environments^3–5^. However, species communities stabilized through antibiotic interactions are not resilient to a high level of spatial mixing^6,7^. Recently, we showed that adding species that degrade antibiotics can lead to communities that are ecologically stable and resilient to spatial mixing^1^. Our model considered species interacting on a grid and consists of the following steps: production of antibiotics around antibiotic-producing species (within area of size *K*_*P*_, radius *r*_*production*_), removal of antibiotics near antibiotic-degrading species (within area of size *K*_*D*_, radius *r*_*degradation*_), killing of antibiotic-sensitive species within the antibiotic zones, and colonization of empty regions by randomly choosing surviving species within a given radius *r*_*dispersal*_ to populate randomly selected sites on the grid of the next generation (Extended Data Fig. 3 in Kelsic et al^1^). In Kelsic et al^1^, we have focused primarily on an analytical form of the model which we call the “mixed inhibition-zone” model, in which we take the limit of complete mixing by taking the dispersal radius to infinity (*r*_*dispersal*_^→∞^). This assumption allowed us to find an analytical solution that was convenient for studying ecological stability, demonstrating community resilient to spatial mixing, species abundance perturbations, the interaction strengths *K*_*P*_ and *K*_*D*_, and differential species growth rates. Communities in the mixed inhibition-zone model are also resilient to invasion by degradation cheaters – mutants that cease degrading a given antibiotic^1^. However, following a personal communication from András Szilágyi, Gergely Boza and István Scheuring, we clarified that this simplifying approximation makes the model sensitive to invasion by production cheaters – mutants that stop producing the antibiotic^2^. Under the mixed-inhibition zone model, a producer cheater can invade the community and lead to its collapse as cells are mixed before they get the self-benefit from killing neighboring species (these cheaters may not take over the community^1^, but can lead to transient dynamics that results in the community collapsing through takeover by another species^2^). We noted therefore that a more biologically realistic implementation of the limit of complete mixing, in which mixing occurs only after the short-range dispersal step (Fig. 1), would be more resilient to production cheaters^2^. This can be understood quite intuitively as antibiotic killing happens preferentially near the producers, giving them an advantage relative to cheaters in colonizing the newly voided spaces prior to mixing.

**Figure 1.**
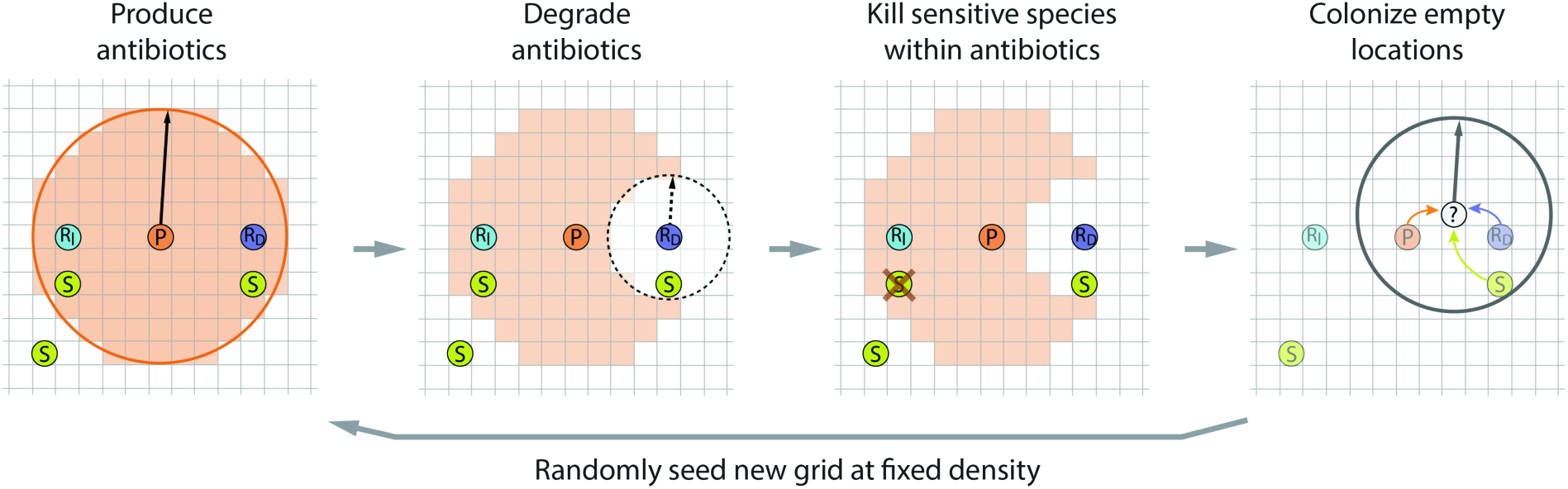
Illustration of the model. A single individual occupies each location on a grid. During each generation: (1) Producer species *P* make antibiotics within radius *r*_*production*_. (2) Antibiotic degrading species *R*_*D*_ remove antibiotics within radius *r*_*degradation*_, for each of the antibiotics. (3) Sensitive species *S* are killed at any locations where antibiotics remain on the grid. Producers and intrinsically resistant species *R*_*I*_ are unaffected by antibiotics. (4) Empty locations on the grid are colonized randomly by any of the remaining species within radius *r*_*dispersal*_. (5) Species are randomly seeded on a new grid at a given density (full mixing).

**Figure 2.**
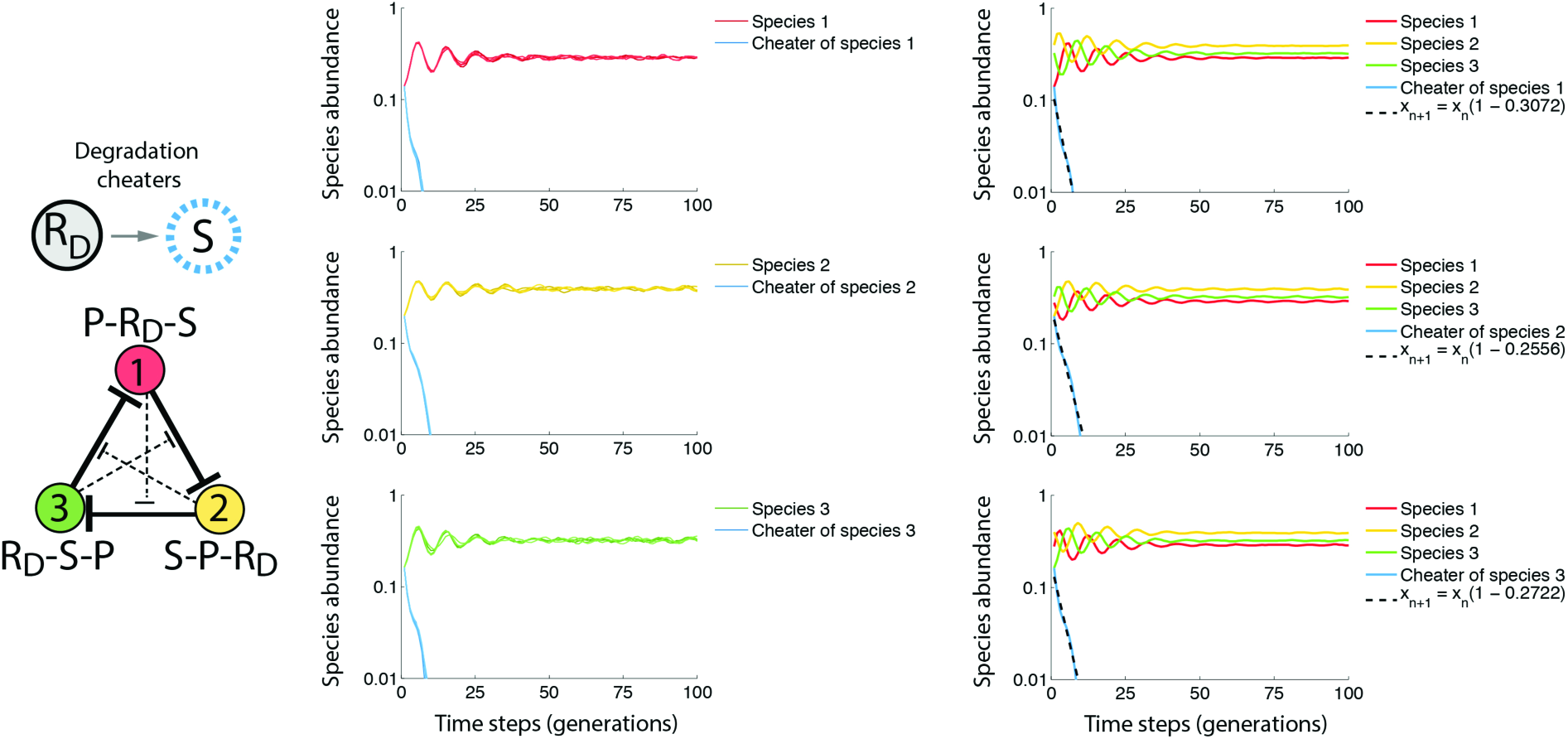
Evolutionary stability of the community against degradation cheaters. We start each simulations of the community at steady state but replace 50% of a given antibiotic degrading species mutated with an antibiotics sensitive species (*D***➔***S*) with equal fitness to its parent. Left plots show parent and cheater traces from 5 different replicas of the simulation. Right plots show the average species abundance of all community members for 25 replica simulations. We estimate cheater fitness (dotted black line) by averaging the rate of exponential decay for the cheater species for each replica simulation until it reached an abundance of less than or equal to 1%. Notably, degradation cheaters drop below 1% in abundance within just a few generations (usually < 25).

**Figure 3.**
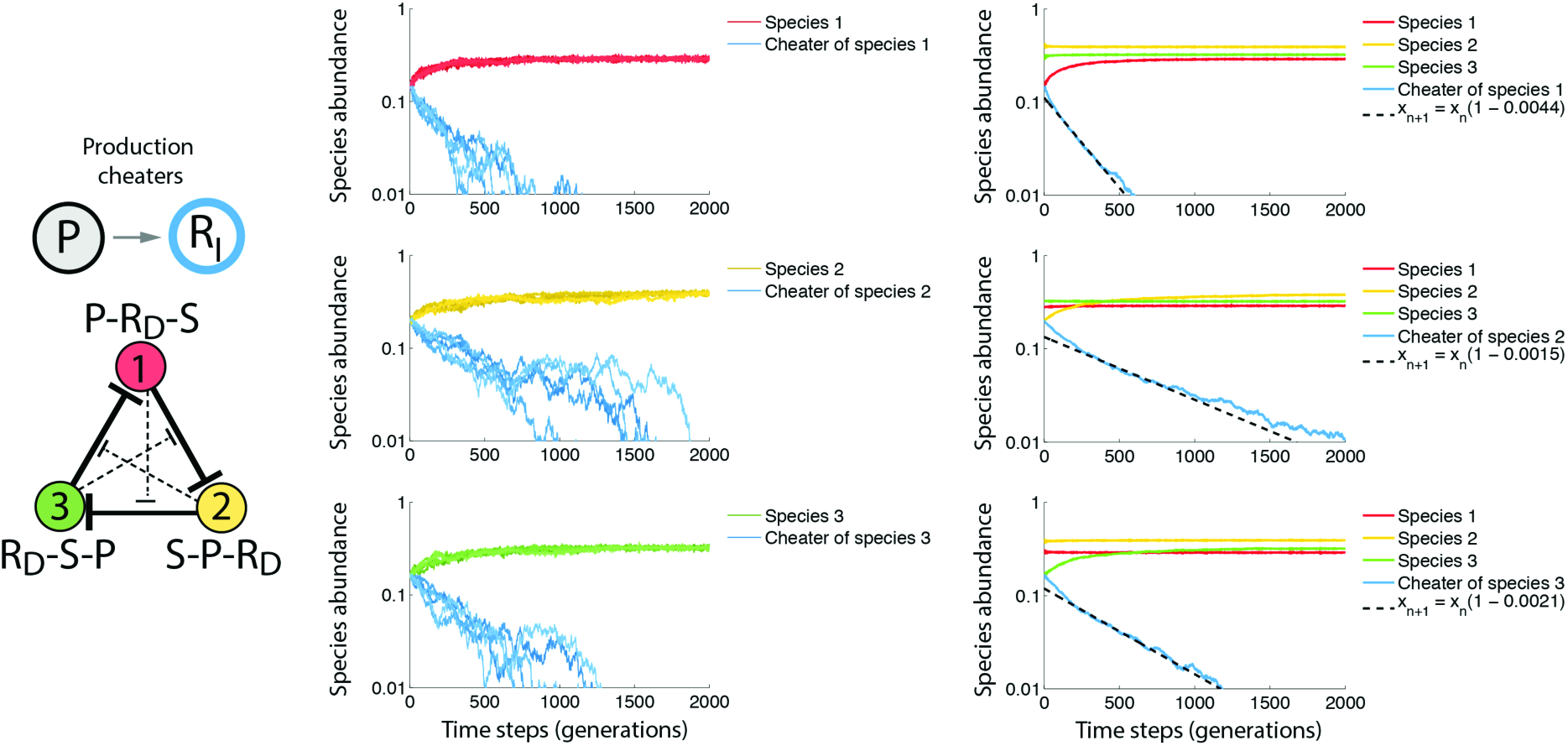
Evolutionary stability of the community against production cheaters. As in Fig. 2 but replacing 50% of a given antibiotic producing species with an antibiotic resistant species (*P***➔***R*_*I*_). Notably, production cheaters are most disadvantaged when their relative abundance is high (steeper slopes, dotted black line shows average of fitness estimates from all 25 simulations until abundance drops below 1%).

To study community resilience to degradation and production cheaters in this more complete model, we simulated the model on a grid with full mixing after the short-range colonization step (Fig. 1). Following Kelsic et al, we considered a cyclic community of three species, setting their inherent growth rates to *g*_1_=1.0, *g*_2_=1.1, *g*_3_=1.2, and the interaction strength to *K*_P_=16 and *K*_D_=3.6 (implemented on a 700×700 grid with density 0.0525, it follows that *r*_*pduction*_=10 and *r*_*degradation*_=5. We set *r*_*dispersal*_=*r*_*production*_). We first allow the community of these three species to reach steady state. Then we introduce either production or degradation cheater for any one of the three species. Cheaters are introduced with intrinsic growth rates equal to that of their parents so that their fitness difference can be assessed from the change in their abundance over time. To prevent the loss of cheater species merely from genetic drift, we study the more stringent situation in which half of the parent species population is converted into the cheater phenotype.

Consistent with our expectation, simulation results show that the community is robust to both production and degradation cheaters. For each run of the simulation, we observe consistently that cheaters decrease in abundance while their parents and the two other species return to their original steady state value (Fig. 2–3). The abundance of all cheaters decreased roughly exponentially until ultimate extinction. We estimated the fitness disadvantage of cheaters relative to their parents by fitting their change in abundance to a decaying exponential and averaging the rate of exponential decay over 25 replicate simulations. As in the mixed-inhibition zone approximation, we find strong selection against degradation cheaters (where *x*_*n+1*_ = *x*_*n*_(*1+s*), *s*_*cheater*_≤ –25%; Fig. 2). Unlike the sensitivity of the mixedinhibition zone approximation to production cheaters^2^, we find that in the complete model production cheaters of all species are selected against with small yet significant selection coefficients (*s*_*1*_= –0.44% ± 0.04%, *s*_*2*_= –0.15% ± 0.02%, *s*_*3*_= –0.21% ± 0.02; deviations are one standard error of the mean, Fig. 3). As expected, selection against production cheaters depends on the radius of production and dispersal, and is maximized when antibiotic production or dispersal occur over smaller distances (Fig. 4).

**Figure 4.**
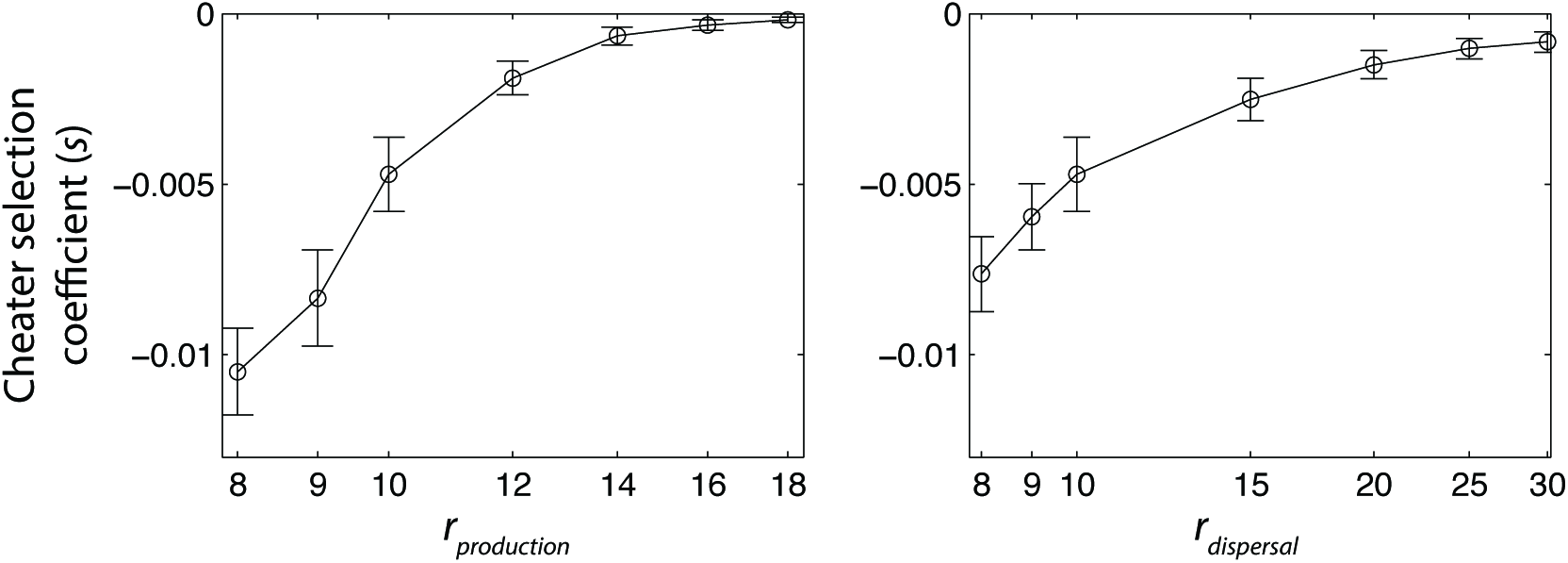
Selection against production cheater is maximized for short-range antibiotic production and colonization. Effective cheater selection coefficients *s*, where *X*_*n+1*_ = *X*_*n*_(*1+s*), for the species 1 production cheater as estimated from averaging rates of exponential decay across 25 replicas, varying *r*_*production*_ (left, fixing *r*_*degradation*_=5 and *r*dispersal =10) or varying *r*_*dispersal*_ (right, fixing *r*_*degradation*_=5 and *r*_*production*_=10). Error bars are two standard deviations from the mean.

To test the generality of selection against production cheaters, we extended our analysis to communities with larger number of species. We identified stable 4-species network topologies by screening random communities in the mixed-inhibition zone model as described in Kelsic et al^1^ (*g*_1_=1.0, *g*_2_=1+0.1/3, *g*_3_=1+0.2/3, *g*_4_=1.1, and interaction strengths *K*_*P*_=7.25 and *K*_*D*_=5.45). We narrowed this set to communities that were also stable in the complete model by running for 500 time steps and confirming no loss of species (implemented on a 500×500 grid with density 0.05, thus *r*_*dispersal*_=*r*_*production*_=7 and *r*_*degradation*_=6). We then tested for selection against production cheaters: for each antibiotic produced by a species in the community, we started 30 replica simulations simulations with 50% of this parent species converted into a production cheater with intrinsic growth rate equal to the parent (Fig. 5). We calculated selection coefficients by calculating the rate of exponential decay of the cheater/parent ratio for each replica and then averaging this value across all replicas. Selection coefficients for production cheaters from 75 randomly selected ecologically stable communities were almost always negative (Fig. 5c; 249 of 251 cheaters had negative selection coefficients, while the remaining 2 cheaters had selection coefficients that were statistically indistinguishable from zero, |s|<stderr). Therefore selection against production cheaters is maintained within the vast majority of these non-canonical network topologies and appears to be general property of such communities.

**Figure 5.**
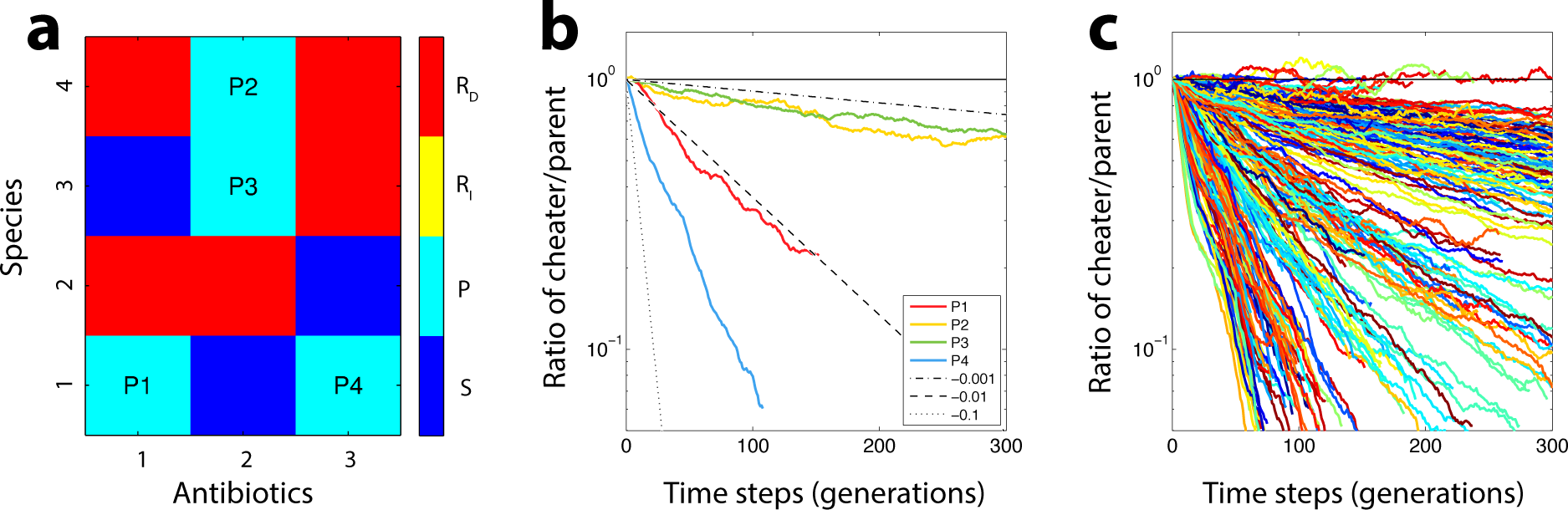
Figure 5. Selection against production cheaters in non-canonical 4 species communities. (**a**) Species phenotypes for an example stable non-canonical 4 species network with possible production cheaters as indicated (P1-P4). (**b**) Ratio of cheater/parent abundances for this example network when starting cheaters at 50% of parent abundance. Cheater/parent ratio was calculated as the average over 30 replica simulations for all time steps up until the first species extinction. Dotted lines show expected slope of selection coefficients ‐0.1, ‐0.01 and ‐0.001. (**c**) Ratios of cheater/parent for all 251 production cheaters from 75 randomly selected stable networks. All cheaters had selection coefficients *s<0* except for 2 cheaters, which were statistically indistinguishable from zero based on the standard error across all 30 replicas (|s|<stderr).

In conclusion, introducing antibiotic degrading species allows the existence of communities resilient to both ecological perturbations and evolutionary loss-of-function mutations. Antibiotic degrading species that lose their ability to degrade become sensitive to the antibiotic and are thereby inhibited more than their parents. Antibiotic producing species that lose the ability to produce will stop killing competitor cells in their vicinity and thereby lose the opportunity to colonize the vacant areas created by this killing. Therefore, these loss-of-function mutants cannot take over a community unless these mutations provide a sufficiently large growth rate advantage. While a short-range colonization step after antibiotic killing is important for selection against production cheaters, long-term spatial structure is not required as community stability occurs even with full mixing at the end of each generation. These results extend to almost every randomly selected non-canonical 4 species network tested and it would be interesting to investigate stability in even larger communities. We emphasize though that such investigation and robustness to production cheaters would be more appropriately studied using the complete model as simulated here and not the mixed-inhibition zone approximation where there is no selection against production cheaters. Our results offer a self-consistent hypothesis about the mechanisms that allow coexistence of antibiotic producing and degrading species in nature and can help direct the construction of stable synthetic communities.

